# Host replacement dynamics shape virulence and genome evolution in a plant RNA virus

**DOI:** 10.1101/2025.03.27.645716

**Authors:** Santiago F. Elena, María J. Olmo-Uceda, Silvia Ambrós

## Abstract

We have investigated how the dynamics of host replacement affect the evolution of a plant RNA virus. Specifically, we have examined how sudden *vs*. gradual transitions from more susceptible to less susceptible *Arabidopsis thaliana* genotypes influence turnip mosaic virus’ virulence and population genomic diversity. Our results show that the evolution of virulence and the complexity of the mutant swarm is linked to both the type of host succession and the genetic basis of the host resistance. Faster changes in virulence are observed after sudden transitions, with an evolutionary trend towards more severe symptoms that appear later during infection. Gradual transitions resulted in greater population mutational load and higher genetic polymorphism compared to sudden transitions. In contrast, beneficial mutations associated with sudden transitions had stronger fitness effects than those linked to gradual transitions. This research highlights the importance of considering the rate of environmental changes in virus evolution and provides insights into predicting how viruses adapt and evolve in temporally changing environments, with implications for agriculture and public health.

## 1. Introduction

The adaptation of asexual populations to dynamic and complex environments is a central topic in evolutionary biology (Futuyma and Moreno 1988), particularly in the context of emerging infectious diseases (Woolhouse et al. 2001; Bedhomme et al. 2015). While adaptation to constant environments has been extensively studied (Kaltz and Shykoff 1998; Turner and Elena 2000; Kawecki and Ebert 2004), the influence of environmental complexity —specifically, the degree of host genetic diversity for resistance (Schmid-Hempel and Koella 1994; Pfenning 2001; Laine et al. 2014; González et al. 2019; Sallinen et al. 2020) and the rate of environmental change (Turner and Elena 2000; Cuevas et al. 2003; Morley et al. 2015; Morley, Broniewski et al. 2016; Morley, Sistrom et al. 2016)— has received less attention. RNA viruses, with their high mutation rates and ability to rapidly adapt to new hosts (Holmes 2009; Elena et al. 2011; Elena 2016), represent an ideal model system for exploring these evolutionary dynamics. Furthermore, understanding how RNA viruses respond to changes in their environment, particularly in host community composition, is crucial for predicting their potential for emergence and evolution in natural and anthropized environments (Woolhouse et al. 2001; Holmes 2009; Elena et al. 2014; Bonneaud and Longdon 2020).

Previous studies have shown that the rate at which a new host “invades” an environment can significantly impact viral adaptation (Morley et al. 2015; González et al. 2019). In environments where changes occur abruptly, beneficial mutations with large effects tend to become fixed early, followed by mutations with progressively smaller effects. This leads to a specific adaptive trajectory in which early mutations have a greater fitness impact (Bello and Waxman 2006; Collins et al. 2007; Schiffels et al. 2011; Gorter et al. 2016; Morley and Turner 2017). In contrast, in environments with gradual change, populations may converge on similar molecular substitutions and experience stronger clonal interference (Morley and Turner 2017; Somovilla et al. 2019). Additionally, in such environments, the selection coefficients of alleles are time-dependent, altering the dynamics of adaptation.

In the context of ecological succession —the process of change in the species composition of an ecological community over time (Connell and Slatyer 1977)— the dynamics of host replacement from a more susceptible to a more resistant genotype or species mimic natural scenarios where viruses must adapt to new selective pressures. Host ecological succession thus raises a key question: How does the speed of host replacement influence the evolution of virulence and genomic dynamics? Experimental evolution studies conducted in cell cultures with Sindbis virus (SINV) revealed that populations experiencing a gradual transition to a less susceptible host cell type consistently achieved higher fitness levels on the new host and exhibited less variability in genomic outcomes (Morley et al. 2015; Morley and Turner 2017). Moreover, populations exposed to both rapid and gradual changes converged on different genotypes, suggesting a strong influence of historical contingency. These findings underscore the importance of considering the rate of environmental change as a key factor in the evolutionary trajectories of viral populations.

In this study, we explore the pace of host species succession using the experimental evolution pathosystem composed of turnip mosaic virus (TuMV; species *Potyvirus rapae*, genus *Potyvirus*, family *Potyviridae*) and its natural host *Arabidopsis thaliana* (L.) HEYNH (genus *Arabidopsis*, family *Brassicaceae*). Drawing inspiration from Morley et al. (2015), we examine the evolutionary dynamics of TuMV in the context of ecological succession from more to less susceptible *A. thaliana* genotypes over time at different paces. Our primary objective is to determine how the rate of host succession affects virulence, genetic diversity, and phenotypic convergence. Specifically, we compare the outcomes of sudden transitions with those that promote a gradual shift. As a second goal, not addressed by Morley et al. (2015), we evaluate the effect of genetic divergence between reservoir and novel hosts by comparing TuMV evolution under two scenarios: (*i*) host plants differing only in alleles at two well-known resistance-related loci and (*ii*) host plants differing in multiple loci genome-wide, belonging to two distinct ecotypes. Our overarching goal is to explore how different rates of host ecological succession influence viral evolution and adaptive responses. This approach is relevant not only for understanding the evolution of viruses in nature but also for applications in agriculture and healthcare, where controlling the spread of viral diseases requires a deep understanding of their capacity to adapt to new hosts.

## 2. Materials and Methods

### 2.1. Plants and viruses

Two different pairs of plant genotypes were used as hosts in the evolution experiments. The first pair consisted of single mutants of the *JASMONATE INSENSITIVE 1* (*JIN1*) and *ENHANCED DISEASE SUSCEPTIBILITY 8* (*EDS8*) genes, respectively. JIN1 is a negative regulator of the salicylic acid-dependent signaling pathway (Laurie-Berry et al. 2006); the *jin1* mutant has been shown to be highly permissive to TuMV infection (Navarro et al. 2022). In contrast, *eds8-1* overexpresses plant defensin genes and exhibits enhanced systemic acquired resistance (Love et al. 2007), making it less susceptible to TuMV infection (Navarro et al. 2022).

The second pair of hosts comprised of the ecotypes Gy-0 (NASC stock number N78901 from La Minière, France) and Oy-0 (NASC stock number N22658 from Øystese, Norway). González et al. (2019) showed that Gy-0 is significantly more sensitive than Oy-0 to TuMV infection, with Gy-0 also developing milder disease symptoms.

Plants were maintained in a BSL-2 climatic chamber under a photoperiod of 8 h of light (LED tubes at PAR 90−100 μmol/m^2^/s) at 24 °C, followed by 16 h of darkness at 20 °C and 40% relative humidity. The substrate mixture used consisted of 50% DSM WNR1 R73454 substrate (Kekkilä Professional, Vantaa, Finland), 25% grade 3 vermiculite, and 25% 3−6 mm perlite. Pest management was conducted through the introduction of *Stratiolaelaps scimitus* and *Steinernema feltiae* (Koppert Co., Málaga, Spain).

TuMV infectious sap was prepared from TuMV-infected *Nicotiana benthamiana* DOMIN plants, which had been inoculated with the infectious plasmid p35STunos. This plasmid contains a cDNA of the TuMV genome isolate YC5 from calla lily (*Zantedeschia* sp.) (GenBank AF530055.2), under the control of the cauliflower mosaic virus 35S promoter and the *nos* terminator (Chen et al. 2003), as described in previous studies (*e.g*., González et al. 2019). After the infected plants exhibited symptoms, they were pooled and flash-frozen in liquid N_2_. The frozen tissue was homogenized into a fine powder using a Mixer Mill MM400 (Retsch GmbH, Haan, Germany).

For inoculation of *A. thaliana* plants, 0.1 g of the powder was diluted in 1 mL of inoculation buffer [50 mM phosphate buffer (pH 7.0), 3% PEG6000, 10% Carborundum]. A 5 μL aliquot of the inoculum was gently rubbed onto three leaves per plant. All plants were inoculated when they reached growth stage 3.5, according to the Boyes et al. (2001) scale. This synchronization ensured that all plants were at the same phenological stage at the time of inoculation. The initial viral stocks were obtained from the infection of homogeneous populations of the corresponding susceptible plants (passage 0 in Fig. 1).

**Figure 1.**
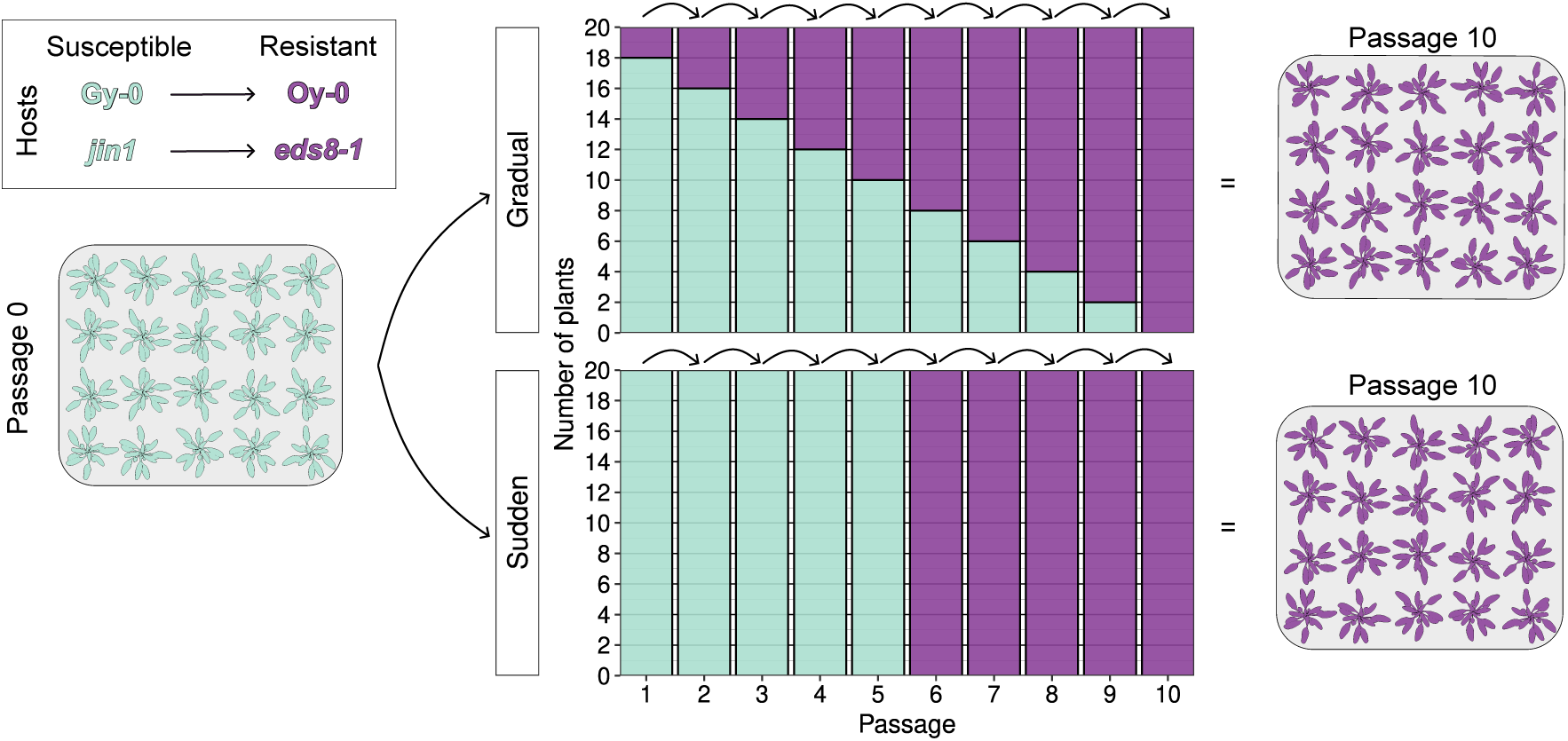
Design of the evolution experiment. Green plants represent the susceptible (reservoir) host, either Gy-0 or *jin1*, while purple plants represent the corresponding resistant (novel) host, Oy-0 or *eds8-1*, respectively. The initial viral stocks were obtained by collecting infected plant populations of the two corresponding susceptible genotypes (passage 0). These infectious saps were then used to initiate the two ecological replacement treatments. In the gradual transition treatment, two resistant plants were added to the population at each passage. In contrast, in the sudden transition treatment, a complete replacement from all susceptible to all resistant plants occurred at passage 6. Each bar represents a plant population with the specified composition (*n* = 20 plants per population).

### 2.2. Experimental evolution design

TuMV lineages were evolved over ten consecutive serial passages, as illustrated in Fig. 1, under two treatments that differed in the pace of host ecological succession. The first treatment referred to as sudden replacement, involved twenty plants of the corresponding susceptible reservoir genotype (*jin1* or Gy-0). Five passages were conducted exclusively in these susceptible plants. Subsequently, all plants were replaced with the novel resistant genotype (*eds8-1* or Oy-0).

The second treatment referred to as gradual replacement, also involved populations of twenty plants. However, with each passage, the number of novel resistant plants was incrementally increased by two, such that by the final passage, all plants in the population were resistant (Fig. 1).

For both treatments, passages were performed by harvesting all plants 12 days post-inoculation (dpi), preparing infectious sap as previously described, and using it to inoculate the next plant population. Three leaves from each 21-day-old plant were rub-inoculated with 5 μL of infectious sap and 10% Carborundum (100 mg/mL).

### 2.3. Phenotyping disease

Three disease-related traits were measured for each plant over 12 dpi: (a) Infectivity, defined as the proportion of symptomatic plants out of 20 inoculated plants at 12 dpi. (b) Area under the disease progress stairs (AUDPS), a single metric summarizing the temporal dynamics of symptom onset (Simko and Piepho 2012); AUDPS ∈ [0, 12]. And (c) average symptom severity at 12 dpi, assessed on a discrete scale ranging from 0 (no infection or asymptomatic infections) to 5 (generalized necrosis and wilting) [see Fig. 1 in Butković et al. (2021)].

### 2.4. RNA extraction and preparation of samples for RNA-seq

Pools were created from 20 symptomatic plants per condition at passages 1, 4, 7, and 10. These were flash-frozen with liquid N_2_ and stored at −80 °C until they were homogenized into a fine powder as in section 2.2. For RNA-seq, total RNA was extracted from 70 to 90 mg of infected and healthy plants using the Plant RNA Purification kit (Agilent Technologies, Santa Clara, CA, USA) according to the manufacturer’s instructions. RNA quality was assessed using a NanoDrop One (Thermo Fisher Scientific, Waltham, MA, USA) and agarose gel electrophoresis, while integrity and purity were evaluated with a Bioanalyzer 2100 (Agilent Technologies). Library preparation and Illumina sequencing were performed by Novogene Europe using a NovaSeq 6000 platform with the Lnc-stranded mRNA-Seq library method, ribosomal RNA depletion, directional library preparation, 150 bp paired-end reads, and 6 Gb of raw data per sample. Novogene Europe performed a quality check of the libraries using a Qubit 4 Fluorometer (Thermo Fisher Scientific), qPCR for quantification, and a Bioanalyzer for size distribution detection.

### 2.5. Bioinformatic methods

The quality of the Fastq files was assessed using FASTQC (Andrews,2010) and MultiQC (Ewels et al. 2016). The files were preprocessed as paired reads with BBDuk (https://sourceforge.net/projects/bbmap/). Adapters were removed, the first ten 5’ nucleotides of each read were trimmed, and sequences at the 3’ end with an average quality below 10 were discarded. Processed reads shorter than 80 nucleotides were excluded. The following parameter values were used: *ktrim* = r, *k* = 31, *mink* = 11, *qtrim* = r, *trimq* = 10, *maq* = 5, *forcetrimleft* = 10, and *minlength* = 80. The processed files were mapped to the TuMV YC5 genome using the BWA-MEM algorithm (Li 2013). The resulting SAM files were binarized and sorted with SAMtools (Danecek et al. 2021), and duplicates were marked using the MarkDuplicates method of GATK version 4.2.2.0 (McKenna et al. 2010).

For TuMV variant calling, viral assemblies without PCR duplicates were used as input for LoFreq (Wilm et al. 2012). Only SNVs with an allele frequency (AF) greater than 5% were used in subsequent analysis, except for selection analyses. Viral load was estimated from the RNA-seq data and expressed as viral reads mapping to the TuMV genome per million bases (VRMB).

The selection coefficient (*s*) and effective population size (*N_e_*) for each variant were estimated using approxwf (Ferrer-Admetlla et al. 2016), which employs a discrete approximation of the Wright-Fisher diffusion model and Bayesian inference to estimate population parameters. The median values of *s* and *N_e_* from their posterior distributions were used in downstream analyses. Confidence intervals were constructed using the Bayesian 95% high posterior densities (HPD) intervals.

### 2.6. Statistical analyses

The time series for each of the three disease-related traits, as well as for the viral load estimated for each evolving population, were fitted to the general autoregressive integrated moving average ARIMA(*p*, *d*, *q*) model:

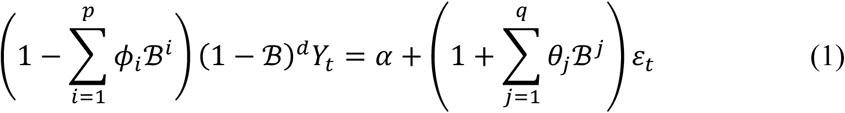

where *Y_t_* represents the estimated value of the corresponding trait at passage *t*; ℬ is the backward shift operator such that ℬ*^n^Y_t_* = *Y_t-n_*; *ϕ_i_* are the coefficients of the autoregressive component (dependence on past values) of order *p* ≥ 0 (number of time lags); *θ^i^* are the coefficients of the moving average component (dependence on past errors) of order *q* ≥ 0; *d* ≥ 0 is the differencing order, representing the number of times the time series is differenced to achieve stationarity; and *ε_t_* is the random error at time *t*, assumed to be independently and identically distributed as *N*(0, *σ*^2^). The coefficient *α* introduces a deterministic component. For example, if *d* = 0, *α* is a constant intercept term; if *d* = 1, *α* introduces a linear trend to the original series; if *d* = 2, *α* introduces a quadratic trend, and so on. Thus, Eq. 1 captures both stochastic components (through *ϕ_i_*, *θ_j_* and *ε_t_*) and deterministic trends (via *α*). ARIMA fittings and model selection were performed using the R package “forecast” version 8.23.0 with automatic estimation of the *11* parameter of the Box-Cox transformation. Four different ARIMA models were significantly fitted to the different time series, as mentioned in the Results section. Firstly, ARIMA(0,0,0) corresponds to a pure random walk with no memory (*i.e*., a mean value with error around it). In this case Eq. 1 reduces to:

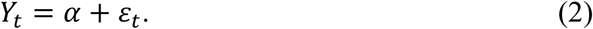

Secondly, ARIMA(1,0,0) corresponds to a stationary process (constant mean and variance over time) with short-term dependency on the mean. In this case, Eq. 1 simplifies to:

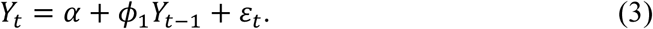

Thirdly, ARIMA(0,1,0) corresponds to a random walk with a temporal tendency. In this case, Eq. 1 reduces to:

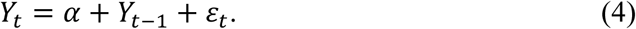

Finally, ARIMA(1,0,1) represents the more complex scenario, where the process has a stationary mean and variance with short-term dependencies on both the mean and the error. In this case, Eq. 1 reduces to the:

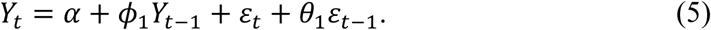

To evaluate the effect of different ecological succession types and genetic architectures of resistance on the evolution of the measured traits, time series were compared pairwise using the dynamic time warping (DTW) algorithm. This method measures the similarity between two temporal sequences (Zhou and de la Torre 2016) as implemented in the R package “dtw” version 1.23-1. The statistical significance of the estimated DTW distance values was evaluated by bootstrapping, with replacement, the values of both curves at the same time points (10,000 pseudosamples).

Partial correlations between traits, controlling for different factors, were computed using the R package “ppcor” version 1.1. Skewness and kurtosis of allele frequency distributions were computed using the R package “moments” version 0.14.1. Wilcoxon signed-rank and Kruskal-Wallis rank sum tests were performed using the corresponding R base functions.

All the above analyses, as well as any other cited in this text, were performed using R version 4.4.2 in RStudio version 2024.12.0+467.

## 3. Results

### 3.1. Rates of evolution for disease-related phenotypes depend on the type of ecological succession and host genetic architecture transitions

Not all the disease-related traits showed a consistent evolutionary trend. Infectivity did not significantly change over the passages, regardless the type of host succession or the genetic architecture of resistance (Fig. 2). In all four cases, the ARIMA(0,0,0) (Eq. 2), representing random noise around a mean, was the best-fitting model. However, after controlling for the effects of ecological succession type and host genetic architecture transitions, a significant correlation was found between infectivity and AUDPS (partial correlation coefficient: *r_p_* = 0.808, 12 df, *P* < 0.001).

**Figure 2.**
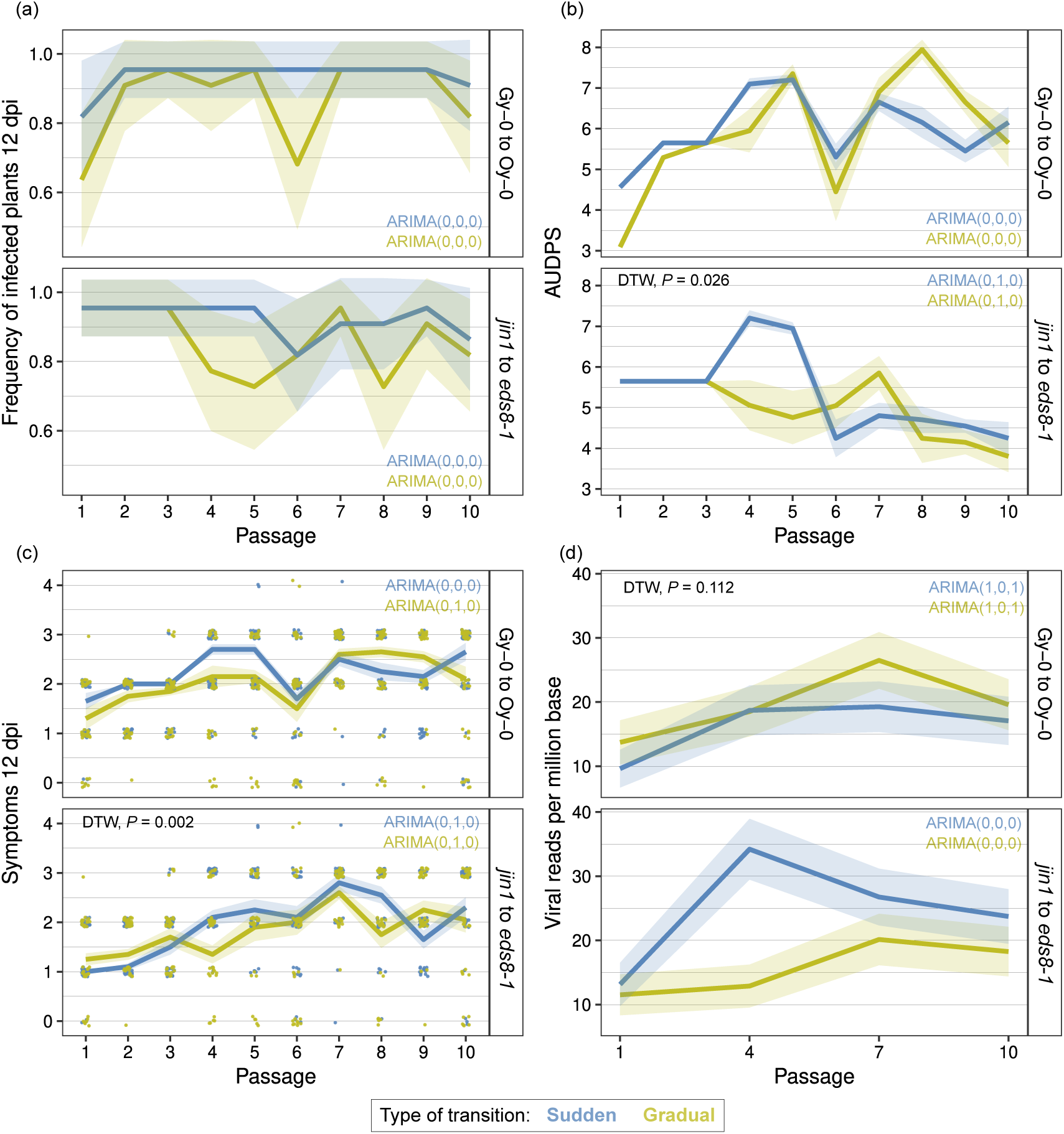
Evolution of disease-related traits and viral load across passages. (a) Infectivity, (b) AUDPS, (c) Average symptoms severity at 12 dpi, and (d) viral load (VRMB). The type of host transition is indicated by different colored lines (blue for sudden, green for gradual), while the two genetic architectures, representing differences between reservoir and novel hosts, are shown in parallel panels (Gy-0 to Oy-0 and *jin*1 to *eds8-1*). Error bands represent ±1 SEM.

By contrast, the evolution of AUDPS was strongly dependent on the genetic architecture of resistance to infection. On the one hand, differences in the succession between the two ecotypes had no overall effect on AUDPS, regardless of the timing of ecological succession [Fig. 2b; best fit to ARIMA(0,0,0) shown in Eq. 2]. On the other hand, the change in AUDPS associated with the ecological succession from the sensitive mutant genotype *jin1* to the more resistant *eds8-1* showed a clear trend [best fit to ARIMA(0,1,0) shown in Eq. 4]. This change was faster in the case of a sudden succession [*α* = −0.331 ±0.078 (mean ± 1 SE), *z* = 4.263, *P* < 0.001] than in the case of a gradual transition (*α* = −0.305 ±0.053, *z* = 5.784, *P* < 0.001), with the difference between the two time series being significant (DTW, *P* = 0.026). The negative value of the deterministic component in both time series indicates that the time required for plants to show visible symptoms increased over the experimental passages. The gradual transition (Fig. 2b, green line) shows a steadier and smoother reduction in AUDPS across passages, whereas the sudden transition from *jin1* to *eds8-1* corresponds to an increase in AUDPS between passages 1 and 4, followed by a sharp decline after all host plants were replaced by the resistant *eds8-1*.

Symptom severity at 12 dpi exhibited a more complex pattern (Fig. 2c). While the sudden transition between the more susceptible Gy-0 and the more resistant Oy-0 ecotypes had no apparent effect [best fit to ARIMA(0,0,0) in Eq. 2], the gradual transition fit better with the ARIMA(0,1,0) model (Eq. 4), exhibiting a significant positive deterministic trend (*α* = 0.106 ±0.039, *z* = 2.724, *P* = 0.003). In the case of the ecological succession from *jin1* to *eds8-1* mutants, both sudden and gradual transitions fit best to the ARIMA(0,1,0) models (Eq. 4), with positive deterministic trends (*α* = 0.138 ±0.051, *z* = 2.725, *P* = 0.003 and *α* = 0.107 ±0.033, *z* = 3.202, *P* = 0.001, respectively). However, the two time series were significantly different (DTW, *P* = 0.002), with the deterministic trend being 29.2% larger in the case of a sudden transition. (Note that the blue line in Fig. 2c increases more rapidly during the first five passages in *jin1* but then declines after the seventh passage. However, for most of the experiment, it remains above the green line corresponding to the gradual transition.)

In summary, disease-related traits exhibit varying evolutionary trends that depend on both the type of host succession and the genetic basis of the resistances involved.

### 3.1. The evolution of viral accumulation depends on the genetic architecture of host transitions

Fig. 2d shows the evolution of viral load (measured as VRMB) as a proxy for TuMV within-host replicative capacity, or absolute fitness. In this case the observed differences were entirely due to the genetic architecture of resistance. Populations undergoing transitions from *jin1* to *eds8-1* mutants showed no significant changes in viral load [best fit to ARIMA(0,0,0)]. However, populations transitioning from the Gy-0 to the Oy-0 ecotypes exhibited significant increases. Both time series best fit the ARIMA(1,0,1) model (Eq. 5), with deterministic trends of *α* = 0.030 ±0.008 (*z* = 3.890, *P* = 0.001) for the sudden transition and *α* = 0.046 ±0.018 (*z* = 2.591, *P* = 0.005) for the gradual transition. However, the difference between the shapes of the two time series was not significant (DTW, *P* = 0.112).

Interestingly, viral load was significantly associated with mean symptoms severity at 12 dpi (*r_p_* = 0.707, 12 df, *P* = 0.005) but not with the other disease-related traits. Fig. 3a illustrates how the evolution of viral load depends on genotype architecture in the context of each phenotypic trait. Symptom severity, a virulence-related trait, increased in all four scenarios; however, a significant increase in viral load was only observed in the transition between the two natural ecotypes. In sharp contrast, infectivity and AUDPS, both transmission-related traits, exhibited opposite trends depending on genotype architecture. A gradual host replacement led to increased transmissibility only when the transition occurred between the two natural ecotypes. Conversely, transitions from susceptible to resistant hosts had a negative impact on infectivity traits, regardless of the replacement pace. Fig. 3b provides a summary of these findings.

**Figure 3.**
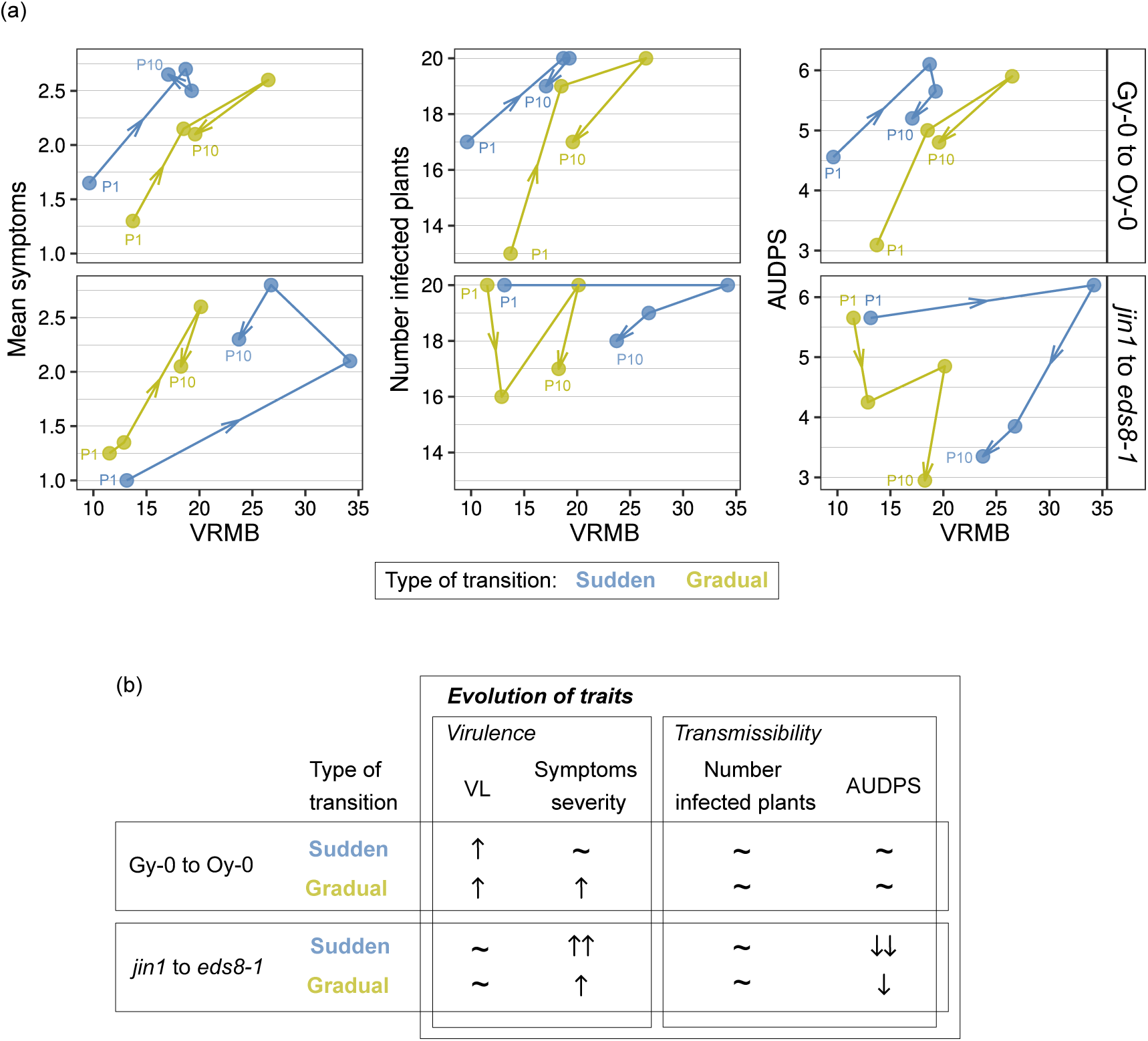
Trajectories of the disease-related traits as a function of changes in viral load. (a) From left to right: mean symptom severity at 12 dpi, infectivity and AUDPS are plotted as a function of viral load (VRMB). The type of host transition is indicated by different colored lines (blue for sudden, green for gradual), while the two genetic architectures, representing differences between reservoir and novel hosts, are shown in parallel panels (Gy-0 to Oy-0 and *jin1* to *eds8-1*). Evolutionary trajectories from passages 1, 4, 7, and 10 are shown in the direction of the arrows. (b) Summary table showing changes in the virulence-related trait (symptoms severity at 12 dpi) and the transmission-related traits (number of infected plants and AUDPS). Arrows indicate whether the evolutionary trend was significant and the direction of change. Double arrows indicate that the DTW test detected a significant difference between the time series being compared. Tildes indicate that the observed changes were better explained by a random walk.

### 3.3. A pervasive trend to increase population mutational load

Fig. 4a illustrates the increase in mutational richness, measured as the number of different mutations with AF > 0.05 observed in each sample, in populations undergoing a sudden host replacement. Different ARIMA models best described the genetic architectures of the host transition. For the transition from Gy-0 to Oy-0 ecotypes, the ARIMA(1,0,1) model (Eq. 5) provided the best fit, with a deterministic trend of *α* = 4.600 ±0.622 (*z* = 7.398, *P* < 0.001). In contrast, the ARIMA(0,0,0) model was more appropriate for the transition from mutant *jin1* to mutant *eds8-1*, with a weaker deterministic trend of *α* = 1.500 ±0.757 (*z* = 1.983, *P* = 0.024), suggesting that the number of different mutations in the first case increased 3.1-fold faster. However, this effect was not significant (DTW, *P* = 0.133) due to noise in the time series.

**Figure 4.**
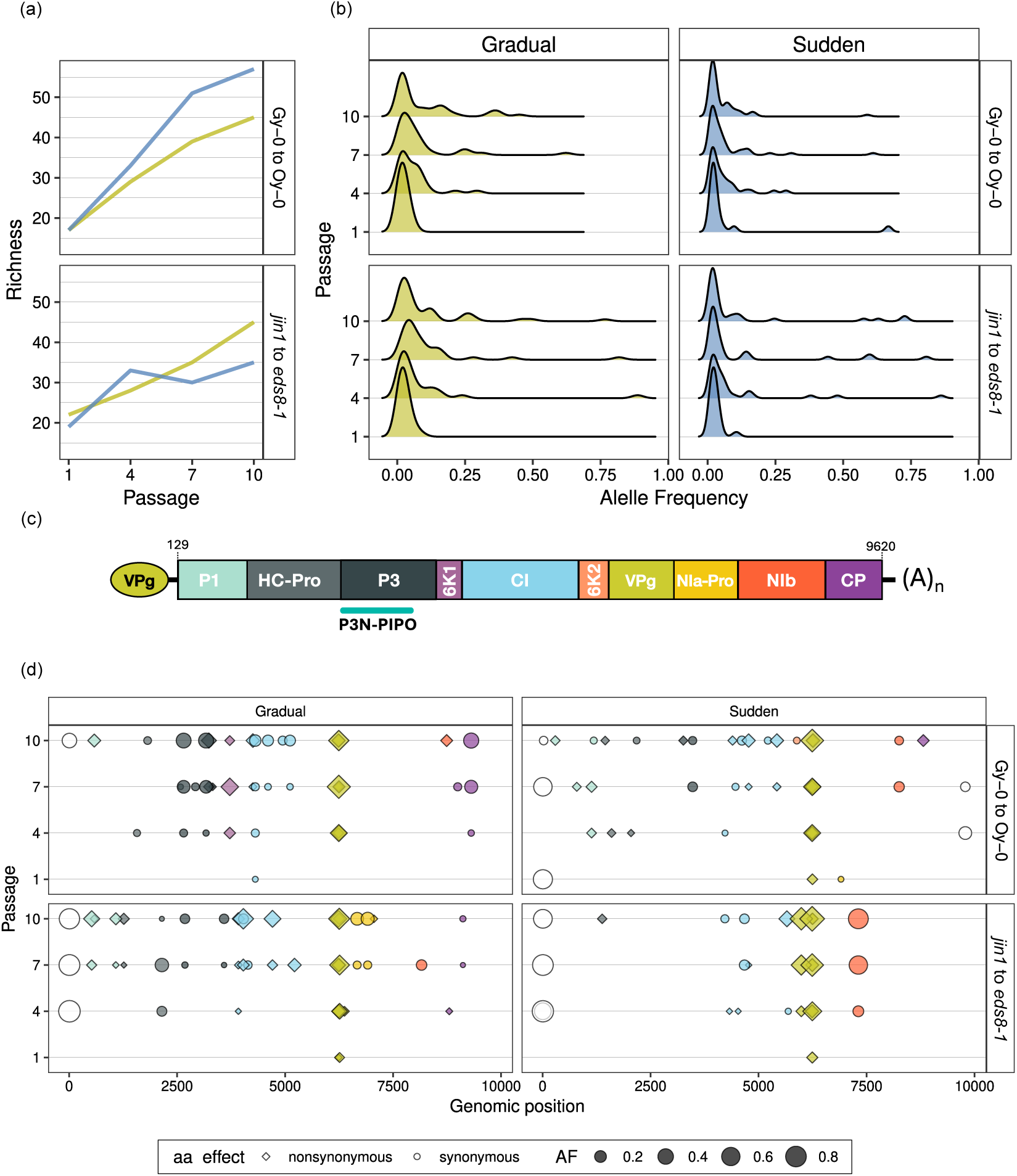
Genotypic changes observed in the different treatments. (a) Mutational richness, measured as the number of different mutations at each passage, for sudden (blue) and gradual (green) host successions. (b) Allele frequency density distributions for transmitted SNVs across different host succession types and genetic architectures. (c) Schematic representation of TuMV genome. (d) SNVs with AF > 0.05. The color of each symbol corresponds to the cistron in which the mutation was observed [as in panel (c)]. Open circles indicate SNVs observed in the 5’UTR. The symbol size is proportional to AF, with circles indicating synonymous mutations and diamonds indicating nonsynonymous mutations.

Fig. 4a also depicts the increase in mutational richness in populations undergoing a gradual host replacement. Here, the ARIMA(0,0,0) model was the best fit for both genetic architectures, with deterministic trends of *α* = 3.133 ±0.320 (*z* = 9.801, *P* < 0.001) for the ecotype transition and *α* = 2.533 ±0.216 (*z* = 11.727, *P* < 0.001) for the mutant transition. A significant difference was observed between the shape of both time series (DTW, *P* = 0.007), with mutational richness increasing 23.7% faster in the gradual replacement of ecotypes compared to mutants.

Interestingly, the sudden transition from Gy-0 to Oy-0 was associated with a 46.8% faster increase in mutational richness (DTW, *P* = 0.035) compared to populations undergoing gradual transitions between the two ecotypes. However, this effect was not observed when comparing transitions between plant genotypes differing in point mutations (DTW, *P* = 0.432).

Fig. 4b presents AF distributions. In all cases, distributions were positively skewed (*g*_1_ > 0) and leptokurtic (*g*_2_ > 3), with heavy tails and sharper peaks than expected under a Gaussian null model. Interestingly, the shape of the distributions changed along passages in a transition-mode-dependent manner. In the case of gradual transitions, distributions showed a significant trend toward widening over time (Kruskal-Wallis test: *P* = 0.017 for Gy-0 to Oy-0 and *P* < 0.001 for *jin1* to *eds8-1*). Specifically, the IQR increased 7.6-fold from passage 1 (0.018) to passage 10 (0.137) in the Gy-0 to Oy-0 transition and 5.2-fold from 0.021 to 0.107 in the *jin1* to *eds8-1* transition. However, in the case of sudden transitions, no significant trend was found (Kruskal-Wallis test: *P* = 0.779 for Gy-0 to Oy-0 and *P* = 0.339 for *jin1* to *eds8-1*).

Fig. 4d shows the distribution of mutations across cistrons, categorized by AF and whether they were synonymous or nonsynonymous. Mutations were more sparsely distributed in the gradual transition, particularly in mutant genotype transitions. Overall, nonsynonymous mutations were more frequent than synonymous changes (Fig. 5a) regardless of treatment or genotic architecture of host resistance (Wilcoxon signed-rank test: *P* σ; 0.021 in all comparisons). However, this pattern was driven largely by the significant excess of nonsynonymous mutations in *VPg* (Figs. 4d and 5b; Wilcoxon signed-rank test: *P* = 0.011 in all comparisons). In contrast, in all other cistrons, the numbers of nonsynonymous and synonymous changes were similar (Figs. 4d and 5b; Wilcoxon signed-rank test: *P* ≥ 0.059 in all comparisons).

**Figure 5.**
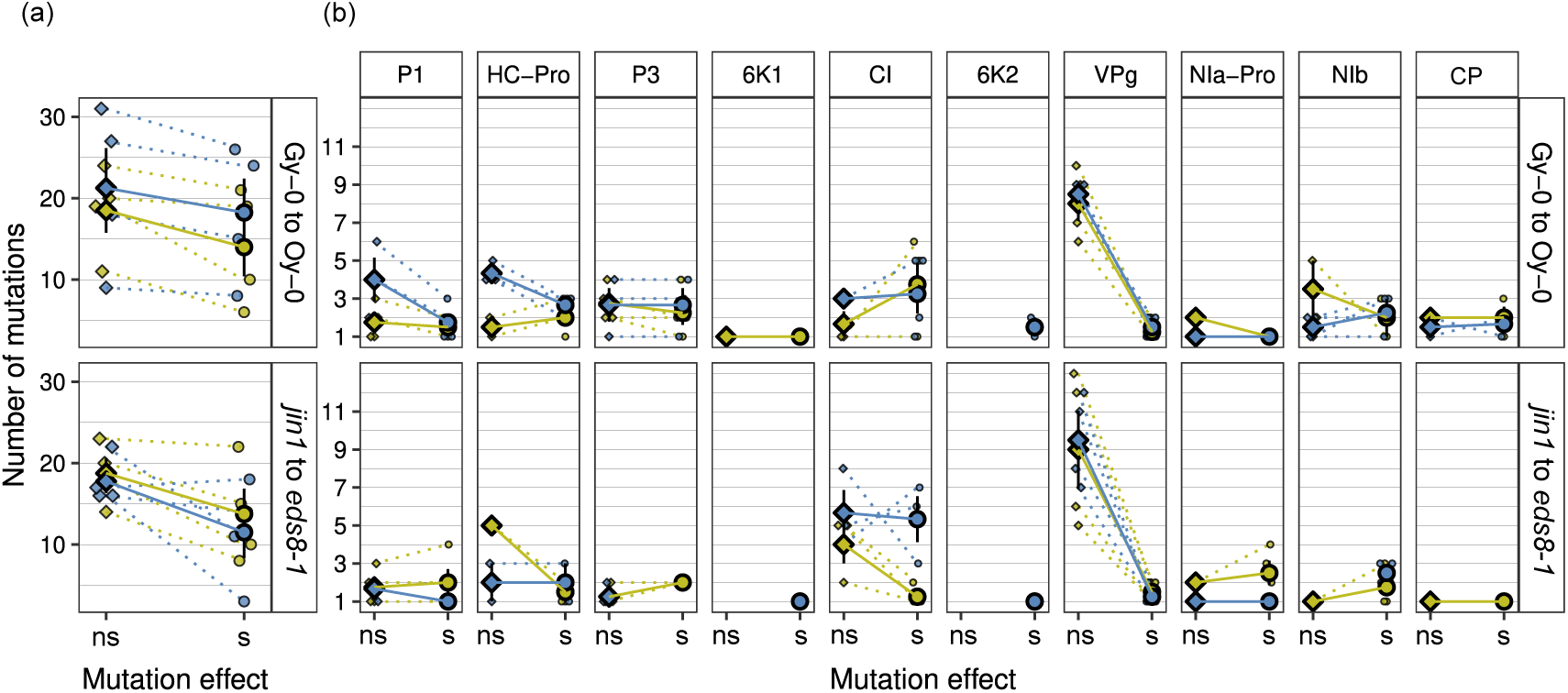
Amino acid effects of the mutations. (a) Total number of nonsynonymous (diamonds) and synonymous (circles) mutations across genotypes for gradual (green) and sudden (blue) host transitions. (b) Number of nonsynonymous (ns) and synonymous (s) mutations per cistron.

This section highlights a pervasive increase in mutational load across all evolving viral populations. However, gradual host transitions led to greater mutational richness and higher genetic polymorphisms (wider allele frequency distributions) than sudden transitions, with the magnitude of these effects influenced by the genetic architecture of host resistance.

### 3.4. Candidate beneficial mutations

Building on the results from the previous section, we evaluated the potential role of selection in shaping the observed mutations. Several mutations persisted across passages (Fig. 4a-b), with some increasing in frequency over time (Fig. 6a). Using a discrete approximation of the Wright-Fisher diffusion model combined with Bayesian inference, we estimated both the selection coefficient (*s*) for each SNV and effective population size (*N_e_*) based on allele frequency changes across passages. Table 1 lists the SNVs with significant selection estimates. For *N_e_*, no significant differences were observed between the two genetic architectures under sudden host transitions. However, under gradual transitions, *N_e_* was significantly smaller in populations transitioning from *jin1* to *eds8-1* than in those transitioning from Gy-0 to Oy-0.

**Figure 6.**
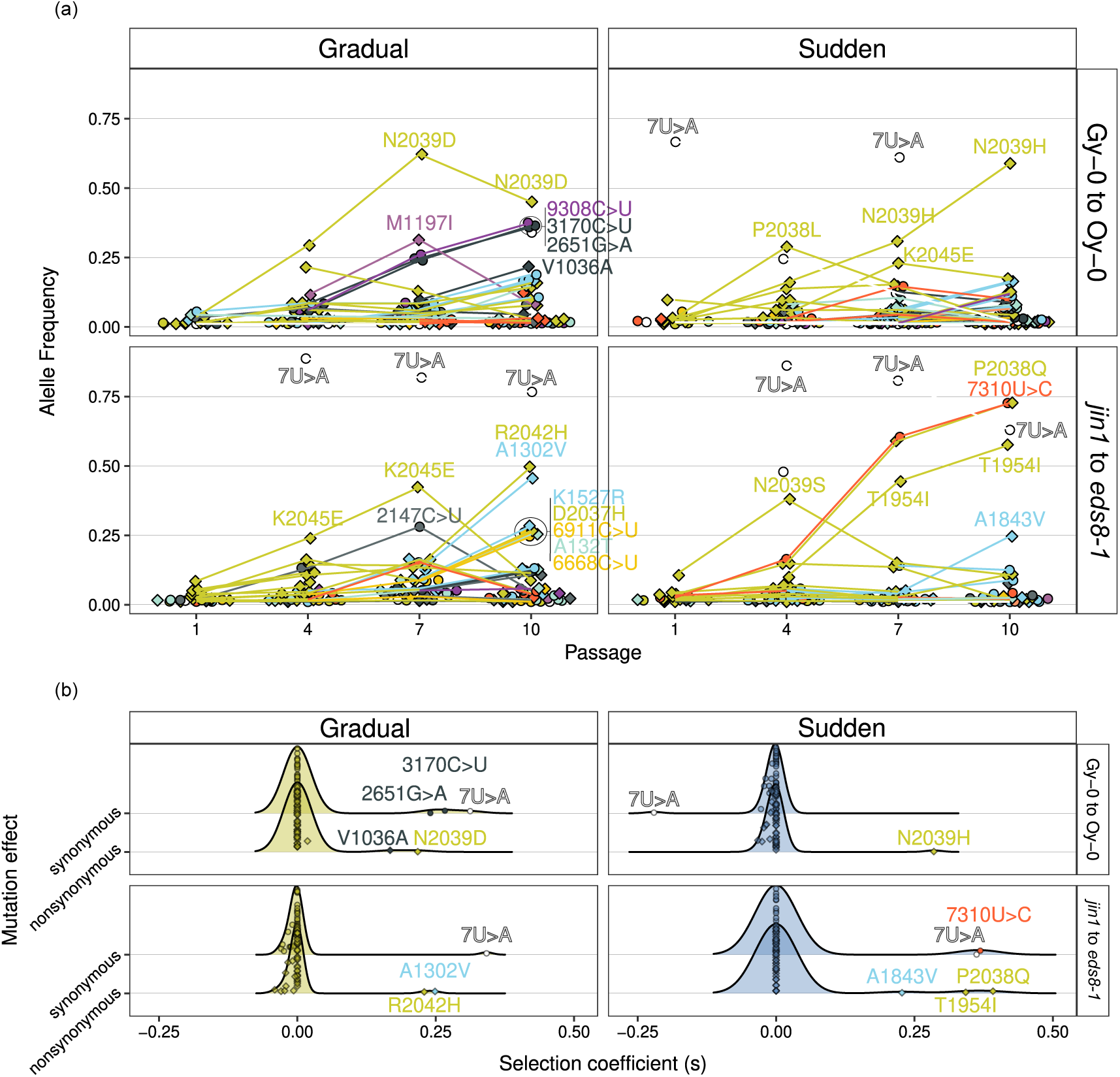
Selected mutations. (a) AF dynamics by treatment and genotype architecture. The shape of dots represents the effect of mutations as in previous plots, nonsynonymous as diamonds and synonymous as circles. Colors by region as in Fig. 4c. (b) Density distribution of the selection coefficients (*s*) per treatment, genotype and mutation effect. Mutations under positive selection are labeled and colored by cistron as in Fig. 4c.

**Table 1.**
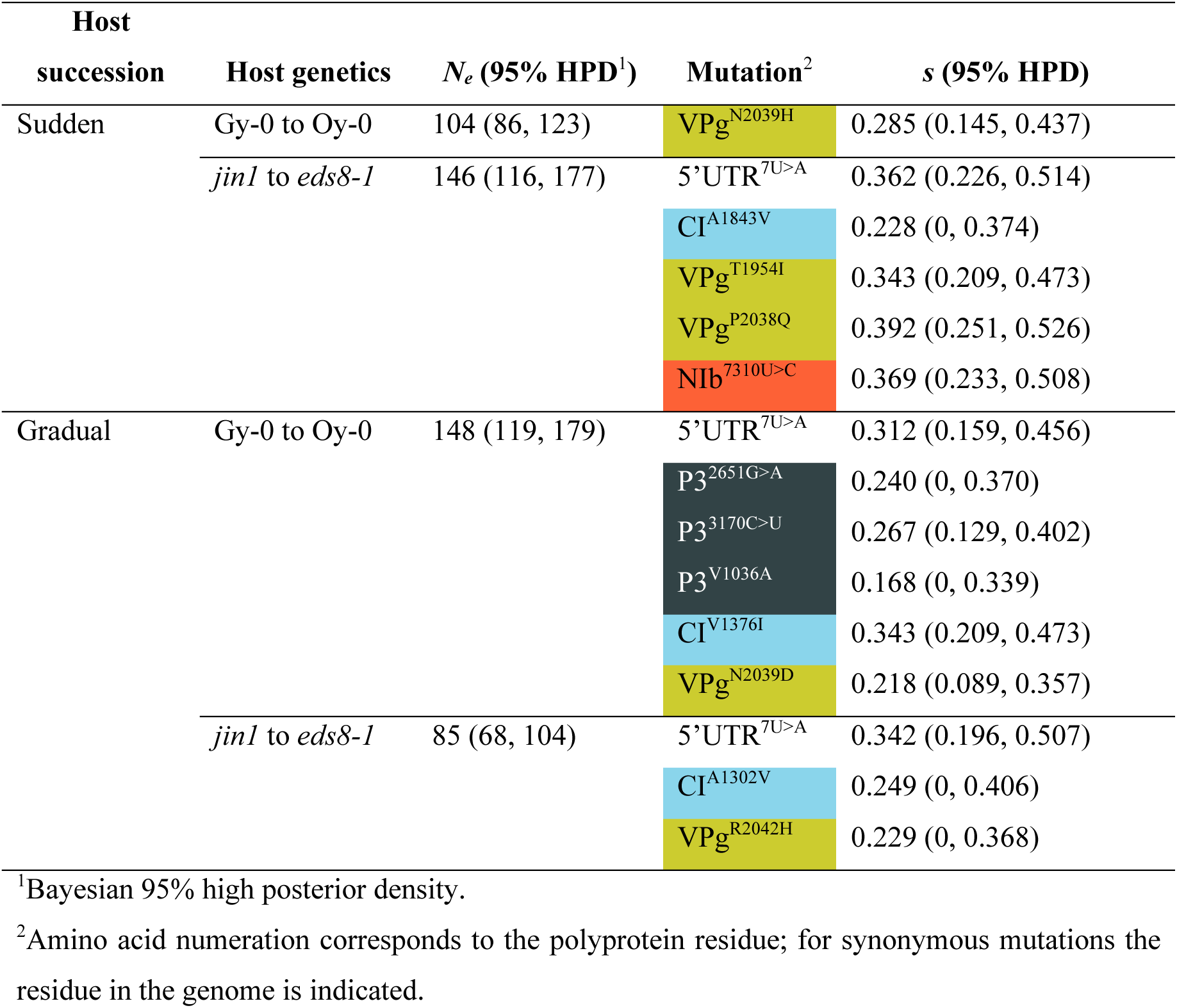
Beneficial mutations identified by the approximated-WF method in each population.

Focusing in the distribution of fitness effects, only positively selected SNVs were identified in the coding regions. Five of these mutations affected VPg, including one polymorphic locus (N2039H and N2039D) observed in two independent populations. Three mutations emerged by passage 3, all in the gradual transition among ecotypes. Additionally, two SNVs were found in CI (both transition schemes for mutant genotypes), and one in NIb (Table 1 and Fig. 6b). Notably, the synonymous mutation NIb^7310U>C^ rose in frequency alongside the nonsynonymous VPg^P2038Q^ replacement during the sudden transition from *jin1* to *eds8-1*, suggesting that the observed fitness value (Table 1) corresponds to that of VPg^P2038Q^, the actual target of selection.

Averaging the fitness effects of candidate beneficial mutations listed in Table 1 revealed that mutations associated with sudden transitions (0.312 ±0.071) are 76.8% stronger than those linked to gradual transitions (0.176 ±0.093), a difference that was statistically significant (*t*_7_ = 2.389, 7 d.f., *P* = 0.048).

As a summary, we identified positively selected SNVs driving adaptation in viral populations, with stronger fitness effects during sudden host transitions compared to gradual ones. Additionally, *N_e_* was influenced by the type of host succession, with significant differences observed under gradual transitions.

## 4. Discussion

In this study, we tracked viral populations evolving in directionally changing heterogeneous hosts, where the progressively novel host acts as a stressor, reducing performance relative to the original host. Under such conditions, the population faces a gradual rather than a sudden host shift and must adapt to a moving optimum as environmental stress intensifies over time (Collins et al. 2007; Koop and Hermisson 2007).

In gradually changing environments, populations experience smaller, incremental drops in fitness and may accumulate a series of mutations with smaller effects (Waxman and Peck 1999; Collins et al. 2007; Koop and Hermisson 2007). The rate of environmental change can influence the likelihood of extinction (Perron et al. 2008; Chevin et al. 2010; Lindsey et al. 2013), fitness levels in the final environment (Lindsey et al. 2013; Morley et al. 2015), and which mutations ultimately become fixed (Waxman and Peck 1999; Koop and Hermisson 2007; Lindsey et al. 2013; Morley and Turner 2017). Consistent with these previous observations, we found that when populations of a plant RNA virus faced the invasion of a novel, more resistant host genotype, the rate of host replacement influenced evolutionary outcomes at both phenotypic and genomic levels.

Specifically, sudden host replacements tended to lead to faster increases in virulence on the less susceptible novel host, although the magnitude of these changes ultimately depended on the complexity of the plant’s genetic resistance architecture. Interestingly, these virulence increases were associated with more severe symptoms, appearing later during infection.

### 4.1. Evolutionary pathways and mutation dynamics

Previous studies have shown that the rate at which a new host “invades” an environment can significantly impact viral adaptation (Morley et al. 2015; González et al. 2019). When host population changes occur abruptly, beneficial mutations with large effects tend to fix early, followed by mutations with progressively smaller effects (Bello and Waxman 2006; Collins et al. 2007; Schiffels et al. 2011; Gorter et al. 2016; Morley and Turner 2017). This results in an adaptive trajectory where early mutations have a greater impact on initial fitness. Conversely, in gradually changing environments, populations may converge on similar molecular substitutions and experience stronger clonal interference (Morley and Turner 2017; Somovilla et al. 2019). Our findings align with this pattern. Regardless of genetic architecture, gradual host transitions were associated with a larger increase in mutational load and more polymorphic viral populations. In contrast, sudden host transitions led to fewer mutations but of larger beneficial effect. Sudden host replacements were also linked to lower genetic diversity, with cases of convergent fixed mutations. Genetic parallelism was common, largely influenced by the genetic architecture of host resistance.

### 4.2. Parallel evolution and key mutations

Notably, we observed amino acid replacements in VPg residue N2039 to H or D specifically during transitions from Gy-0 (susceptible) to Oy-0 (resistant). Similarly, the mutation VPg^R2042H^ was linked to the gradual transition from *jin1* to *eds8-1*. Mutations at these two VPg residues have been identified in prior studies of TuMV evolution in various *A. thaliana* ecotypes and environmental stress conditions (González et al. 2021), plants at different developmental stages (Melero et al. 2023), plants deficient in resistance genes (Navarro et al. 2022), or plants lacking epigenetic regulators (Ambrós et al. 2024).

The mutation VPg^R2042H^ is particularly noteworthy. Carrasco et al. (2024) demonstrated that it exhibits a differential interactome, binding more strongly than the wildtype VPg to several host proteins, including RHD1, involved in DNA methylation; PRCC and PPDK-RP1, regulators of protein modification and signaling; OBE1, which interacts with WRKY transcription factors regulating stress responses; and MYOS3, involved in Golgi, peroxisome, and mitochondrial trafficking.

Additionally, the VPg^P2038Q^ mutation, observed during sudden transitions from *jin1* to *eds8-1*, has been reported previously (Ambrós et al. 2024). Notably, P2038Q, N2039D, and R2042H all occur in a highly variable region of the VPg core domain, which plays a critical role in TuMV adaptation by modifying interactions with the ^m7^G cap-binding pocket of eIF(iso)4E (Carrasco et al. 2024). The reproducibility of these mutations across experiments suggests they enhance YC5 fitness in *A. thaliana* in a genotype-independent manner.

In contrast, mutations such as VPg^T1954I^ and those in P3 and CI have not been reported in previous experimental evolution studies, warranting further investigation.

### 4.3. Comparison with other viral evolution studies

Despite some parallels with prior research, key differences exist. SINV studies found that gradual replacement of susceptible cells led to higher viral fitness (Morley et al. 2015), whereas a Qβ phage study showed that sudden temperature shifts resulted in greater fitness gains (Somovilla et al. 2019). We found no significant effect of host replacement rate on viral accumulation; both treatments reached similar levels. However, viral accumulation was influenced by genetic architecture, with significant fitness increases only in transitions between ecotypes.

SINV (Morley and Turner 2017) and Qβ (Somovilla et al. 2019) studies reported that gradual environmental changes resulted in less variable viral populations. We observed this only in natural ecotypes. In contrast, in *jin1* to *eds8-1* transitions, sudden replacements reduced genomic variability, likely due to a few adaptive mutations sweeping the population.

These discrepancies likely reflect fundamental differences in host systems: SINV and Qβ studies used cell cultures, whereas our study examined multicellular hosts. Replication events per passage are significantly higher in whole plants.

### 4.4. Implications for plant virus management

Our findings have practical implications for plant virus management in agriculture. Gradual crop replacements (*e.g*., introducing resistant varieties over time) may allow pathogens to adapt more slowly, potentially leading to less aggressive strains. Sudden crop replacements (*e.g*., rapid shifts to resistant varieties) create strong selection pressures, driving the emergence of more virulent strains. Sudden replacements may also reduce genetic diversity, favoring a few highly adaptive mutations, while gradual replacements maintain higher genetic diversity, potentially slowing the emergence of resistance-breaking traits. Gradual replacement strategies could enhance resistance gene durability, spreading selection pressure over time, while sudden shifts may accelerate resistance breakdown, leading to economic and environmental costs.

### 4.5. Conclusions

We provide further evidence that the rate of novel host invasion shapes viral adaptation at both phenotypic and genomic levels. This study underscores the need for evolutionary virologists to move beyond simplistic sudden environmental shifts (Elena et al. 2011; Elena 2016) and consider the complexity of natural host communities, which vary across space and time (Woolhouse et al. 2001).

## Acknowledgements

We thank Francisca de la Iglesia for performing the serial passages and for excellent technical support. This work was supported by grants PID2022-136912NB-I00 funded by MCIU/AEI/10.13039/501100011033 and by “ERDF a way of making Europe”, and CIPROM/2022/59 funded by Generalitat Valenciana to SFE. MJOU was supported by grant FPU2019/05246 funded by MCIU/AEI/10.13039/501100011033 and by “ESF investing in your future”.

## Author contributions

SFE conceptualized the idea of the study. SA performed investigation. SFE and MJOU curated the data. SFE, MJOU and SA designed methodology. SFE and MJOU performed formal analysis. SFE wrote the original draft. All authors reviewed and edited the manuscript. SFE acquired funding.

## Data availability

Raw disease-related phenotypic data, with a link to the R code used for the analysis of temporal series, are available at Zenodo repository doi: 10.5281/zenodo.15088312. RNA-seq data can be downloaded from SRA Bioproject code PRJNA1220327. R and bash codes for sequence data analysis and presentation are available in https://gituhub.com/MJmaolu/HostTransitions.

